# Eye–Head Coordination During Rapid Gaze Shifts in Soccer Scanning

**DOI:** 10.1101/2025.09.19.677484

**Authors:** Junki Saito, Tomohiro Kizuka, Seiji Ono

**Author notes:** **Correspondence:** Seiji Ono.

## Abstract

The visual exploratory behavior involving rapid gaze shifts that supports cognitive processes is called “scanning.” Scanning in soccer is a foundational behavior that enables players to explore their environment, supporting rapid and accurate decision-making. However, since most previous studies have focused on head movements, the coordination structure of gaze shifts, including the ocular contribution, is insufficiently understood. This study aimed to determine the coordinated roles of the eye and head during scanning. Twenty male collegiate soccer players performed a passing task paired with video-based situational judgments while eye and head movements were recorded by a 200 Hz sampling rate using an eye tracker system with a built-in gyroscope. We detected eye and head velocities and amplitudes during rapid gaze shifts. As a result, peak eye velocity was significantly higher than peak head velocity (d = 4.09, p < .001). The cross-correlation (CC) between gaze and eye velocities (0.98) was significantly greater than that between gaze and head velocities (0.90). These results indicate that gaze shifts were driven primarily by eye movements rather than head movements. Furthermore, eye and head velocities were negatively correlated (r = −.60, p = .005), whereas their velocity-control profiles (mean velocity normalized by amplitude) were positively correlated (r = .90, p < .001), indicating individual strategy-dependent allocation of effort while preserving control capability of eye and head movements. Our findings provide a characterization of the eye–head coordination structure of gaze shifts in soccer scanning, suggesting the need for comprehensive assessments beyond head-motion metrics alone.

## 1 Introduction

In sports such as soccer, where play rapidly alternates between attack and defense within a wide field of view, the quality of cognitive processing is a critical determinant of performance (Davids et al., 2005; Mann et al., 2007). Visual exploratory behaviors such as gaze shifts involving head movements that support cognitive processes are referred to as “scanning” (Jordet, 2005; McGuckian et al., 2020). It is known that greater frequency and quality of scanning enable players to anticipate more information and gain a competitive advantage. Indeed, observational studies have shown that scanning is a foundational behavior supporting decision-making (Aksum et al., 2021). Furthermore, recent studies at international soccer tournaments have quantified the head direction (Pokolm et al., 2023) and the frequency of scanning (Pokolm et al., 2022), providing the importance of scanning as an essential component for performance.

Previous research on scanning, while emphasizing its importance, has primarily focused on the frequency of head movements and the characteristics of fixations. For example, in soccer, the number and timing of head turns have repeatedly been reported to be associated with performance (Jordet, 2005; McGuckian et al., 2020; Jordet et al., 2020). In addition, the duration and location of fixations are known to influence the accuracy of situational judgments (Roca et al., 2011; Assis et al., 2021). Moreover, visual exploratory behavior has been shown to support tactical decision-making even in youth players (Klatt et al., 2021). These findings provide strong support for the importance of scanning. However, many previous studies have used head turn frequency as a proxy measure of scanning (Aksum et al., 2021; McGuckian et al., 2019). Studies that have simultaneously measured eye movements are extremely rare (Van Biemen & Mann, 2024).

Another study focusing on fixations has emphasized static gaze locations “where one looks” rather than sufficiently addressing the dynamic aspects of gaze shifts and their coordination structure. Recent reviews have also pointed out that although advances in eye-tracking technology have increased laboratory-based studies, knowledge of ocular dynamics under ecologically valid competitive conditions remains lacking (Kredel et al., 2023). Furthermore, even comprehensive reviews addressing the relationship between decision-making and visual exploratory behavior have not explicitly examined the coordination structure of gaze shifts (Silva et al., 2020).

Gaze shifts are a foundational element of scanning, enabling players to efficiently acquire diverse environmental information and providing the prerequisite basis for subsequent decision-making (Roca et al., 2014; Hüttermann et al., 2018). Aksum et al. (2021b) defined in-match scanning as “active head and eye movements performed with the intention of momentarily averting one’s gaze from the ball to search for information about teammates, opponents, referees, or spaces relevant to the unfolding play.” This definition implies that gaze shifts, momentarily diverting the eyes from the ball for information gathering, are indispensable components of scanning. Furthermore, studies in junior and youth players have reported that the efficiency of exploratory patterns directly influences action selection (Klatt et al., 2022; Assis et al., 2021). Exploratory activity, including gaze shifts, has also been demonstrated as a core factor supporting the accuracy and efficiency of decision-making (Mann et al., 2007), with attentional flexibility identified as one of the underlying cognitive abilities (Hüttermann et al., 2018).

In contrast to practical research, basic research has examined the mechanisms of eye–head coordination during gaze shifts in detail, reporting changes in ocular and head contributions as a function of gaze amplitude (Freedman, 2008), individual differences (Fuller, 1992), and physiological constraints of ocular motion (Bahill et al., 1975; Zangemeister et al., 1981). Although there has been a study in the applied sports contexts to reveal characteristics of eye–head coordination in referees’ on-field visual exploration (Van Biemen et al., 2023), investigations probing angular-velocity profiles and qualitative aspects of coordination remain scarce. Therefore, it remains unclear whether the mechanisms of gaze shifts can be applied to the practical context of scanning in soccer.

Based on these findings, the next question in scanning research is not only to examine “where one looks” or “how often the head turns,” but also to elucidate the dynamics of “how the gaze is shifted.” Therefore, the purpose of the present study is to quantitatively clarify the coordination structure of gaze shifts in soccer scanning, specifically examining (1) the comparative dominance of the eyes and the head in forming gaze, (2) the relationships between eyes and head velocities, and (3) their respective functional roles. The findings of this study have the potential to contribute not only to the theoretical advancement of scanning research but also to improved assessment and coaching practices in applied training settings.

## 2 Materials and Methods

### 2.1. Participants

Participants were 20 male university soccer players (mean age: 19.8 ± 1.4 years; height: 171.4 ± 6.7 cm; body mass: 66.8 ± 7.5 kg; playing experience: 13.7 ± 2.8 years). All participants were right-foot dominant, had normal or corrected-to-normal vision, and reported no motor impairments. Many of the participants were regular starters on teams competing in the Kanto University Soccer League and possessed a sufficiently highly competitive level. A priori power analysis using G*Power software (Faul et al., 2007) was conducted to determine the sample size, and the number of participants was set based on the results of this analysis. The study was conducted in accordance with the Declaration of Helsinki (2013), and all experimental protocols were approved by the Research Ethics Committee of the Institute of Health and Sport Sciences, University of Tsukuba. Prior to participation, all participants were informed of the study’s purpose and procedures and provided written informed consent.

### 2.2. Procedure

In this study, we used an experimental task simulating a soccer passing situation to examine the relationship between head and eye movements and gaze-shift strategies during scanning (Fig. 1). Participants stood at a passing location 2.4 m in front of a large screen (280 x 480 cm; Fig. 1-①, ②) and executed a one-touch pass toward a ball (Mikasa FT-550b FPQ) released from a ramp placed 90° to their left (custom-made apparatus: L-shaped angle; Fig. 1-③). Simultaneously with the ball becoming visible, a 2-versus-2 offensive–defensive video was projected onto the screen (EPSON EB-1771W; Fig. 1-④), and participants selected the pass direction toward the side displaying an unmarked teammate. Each participant completed 102 trials, comprising two ball-speed conditions x 17 trials x 3 blocks. After release, the ball was temporarily occluded by a custom-made visual shield (L-shaped angle with plastic board; Fig. 1-⑤) on the ramp and reappeared at a position 3 m before the passing point. An infrared sensor (Arduino Uno R3; Fig. 1-⑥) was placed at the reappearance point, triggering the projection of the situational judgment task on the screen as the ball passed. This setup reproduced contextual features of actual soccer, including “anticipation of pass-impact timing,” “maintenance of ball gaze at the moment of impact” (Van Biemen & Mann, 2024), and “decision-making under time pressure.” Participants were instructed to maintain their gaze straight toward the ramp while the ball was occluded, and then to initiate a gaze shift toward the screen once the ball reappeared, as illustrated in Figure 1.

**Figure 1.**
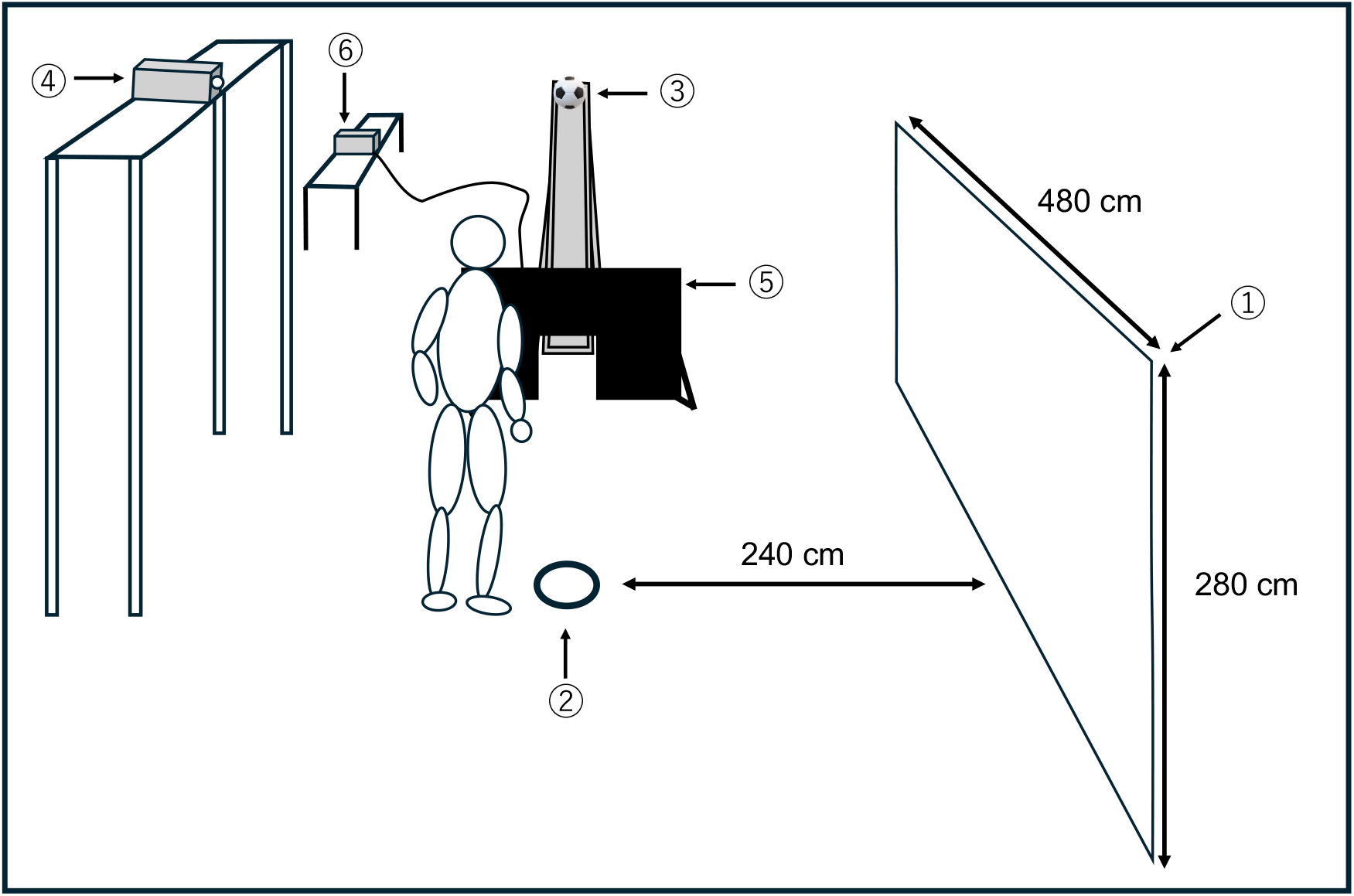
Overview of the experimental setup. (1) Screen; (2) Passing location; (3) Ball-release ramp; (4) Projector; (5) Occlusion screen; (6) Infrared sensor.

### 2.3. Apparatus

Eye movements during the task were recorded using an eye-tracking device (Pupil Invisible glasses; Pupil Labs, Berlin, Germany). The sampling rates of the eye cameras and the scene camera were 200 Hz and 30 Hz, respectively, and eye-position data were acquired at 200 Hz. Head movements were recorded with the device’s built-in gyroscope at a sampling rate of 200 Hz.

### 2.4. Data processing

Pixel values obtained from the eye-tracking device were converted to visual angles based on the device specifications (horizontal: 82°/1088 pixels; vertical: 82°/1080 pixels). The coordinate origin of the scene camera was set at the center of the screen. Eye angular velocity was calculated by differentiating position data, head angular velocity was obtained directly from the gyroscope, and head position data were computed by integrating head velocity. Figure 2A shows a representative example of mean positional waveforms during the task, and Figure 2B presents a representative example of raw positional waveforms. In this study, following prior work, the onset of scanning (Time 0 ms) was defined as the moment when horizontal head angular velocity exceeded 125 deg/s (McGuckian et al., 2019). Data were segmented from −200 ms relative to this onset until the end of each trial. As participants were instructed to align their head and eyes toward the ramp at trial onset, head and eye positions at −200 ms were normalized to 0°. Furthermore, the “gaze-shift phase” was defined as the interval from when gaze angular velocity exceeded 250 deg/s until it fell below this threshold again. This threshold was selected as a conservative criterion for capturing dynamic, match-like movements in this study, based on prior criteria for head movement (125 deg/s; Chalkley et al., 2018; McGuckian et al., 2019), saccade velocity thresholds (120–150 deg/s; Gregori et al., 2016; Kishita et al., 2020), and gaze-shift velocity thresholds (~240 deg/s; Parker et al., 2023; Manakhov et al., 2024). This threshold was intended to robustly capture relatively slow head-driven rotations as well as fast eye-driven movements. Analyses were restricted to horizontal (yaw-axis) movements. Signals were low-pass filtered using a 4th-order Butterworth filter with a cutoff frequency of 50 Hz. The computed metrics included peak velocity, mean velocity, and amplitude for both eyes and head; coefficients of variation (CV); cross-correlations with gaze; and mean-velocity-to-amplitude ratios, as illustrated in Figure 2.

**Figure 2.**
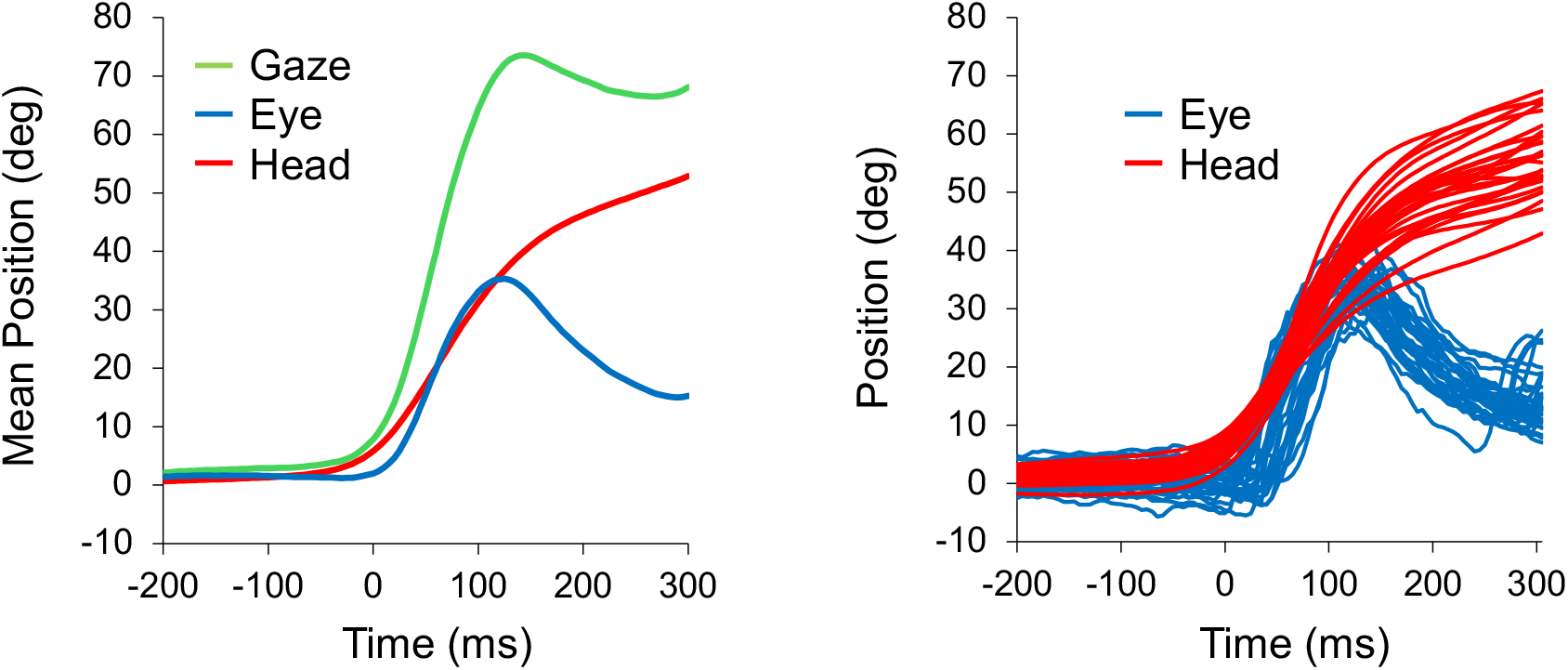
Representative examples of eye and head position traces. (A) Mean waveforms of gaze (green), eyes (blue), and head (red) positions. Position traces are plotted as a function of time, with the trial onset position normalized to 0° for both eye and head traces. (B) Representative positional traces from a single trial. Eye (blue) and head (red) positions are plotted as displacements from 0° at trial onset.

### 2.5. Statistics

Paired t-tests or Wilcoxon signed-rank tests were used to compare the eye and head variables listed in Section 2.4, as well as the time conditions (800 ms vs. 1600 ms). Effect sizes (Cohen’s d or r) were also calculated. Cross-correlation analyses were conducted to examine the relationships between gaze and eye movements and between gaze and head movements (correlation coefficients reported as CC). In addition, Pearson’s correlation analyses were used to examine associations between two variables. Because independent hypotheses were tested for each variable, no corrections for multiple comparisons were applied. For all tests, the significance level was set at α = .05. Statistical analyses were performed using IBM SPSS Statistics version 30 (SPSS Inc.). Unless otherwise noted, data are presented as mean ± standard deviation.

## 3 Results

### 3.1. Comparison between eye and head movements during gaze shifts

Figure 3A illustrates representative examples of peak velocity and amplitude for the eyes and head. As shown in Figure 3B, paired t-tests revealed that peak velocity was significantly higher for the eyes than for the head (t(19) = 18.31, p < .001, d = 4.09). Mean velocity (t(19) = 4.77, p < .001, d = 1.07) and amplitude (t(19) = 5.97, p < .001, d = 1.34) were also significantly greater for the eyes than for the head. Figure 4A presents representative waveforms of mean velocity for gaze, eyes, and head. As shown in Figure 4B, cross-correlation analysis revealed that the maximum correlation coefficient (CC) was 0.98 for gaze–eye and 0.90 for gaze–head, with Wilcoxon signed-rank tests indicating that gaze–eye CC was significantly higher (Z = 3.92, p < .001, r = .88). These results indicate that gaze shifts are led by the eyes, while the head plays a supportive role, as illustrated in Figures 3 and 4.

**Figure 3.**
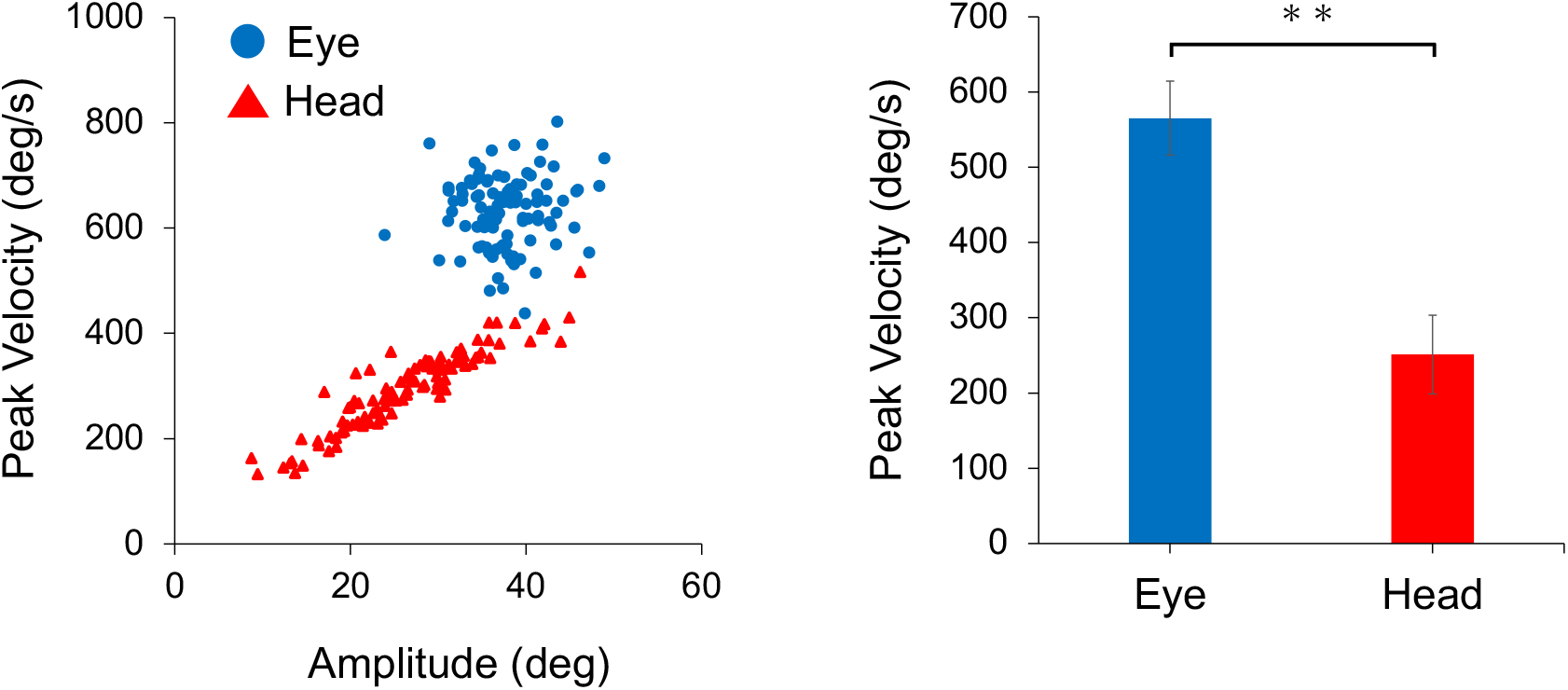
Scatterplots of eye and head peak velocity. (A) Scatterplots of eye (blue) and head (red) peak velocity versus amplitude across trials. (B) Comparison of peak eye and head velocity showing higher eye velocity values (**p < .001).

**Figure 4.**
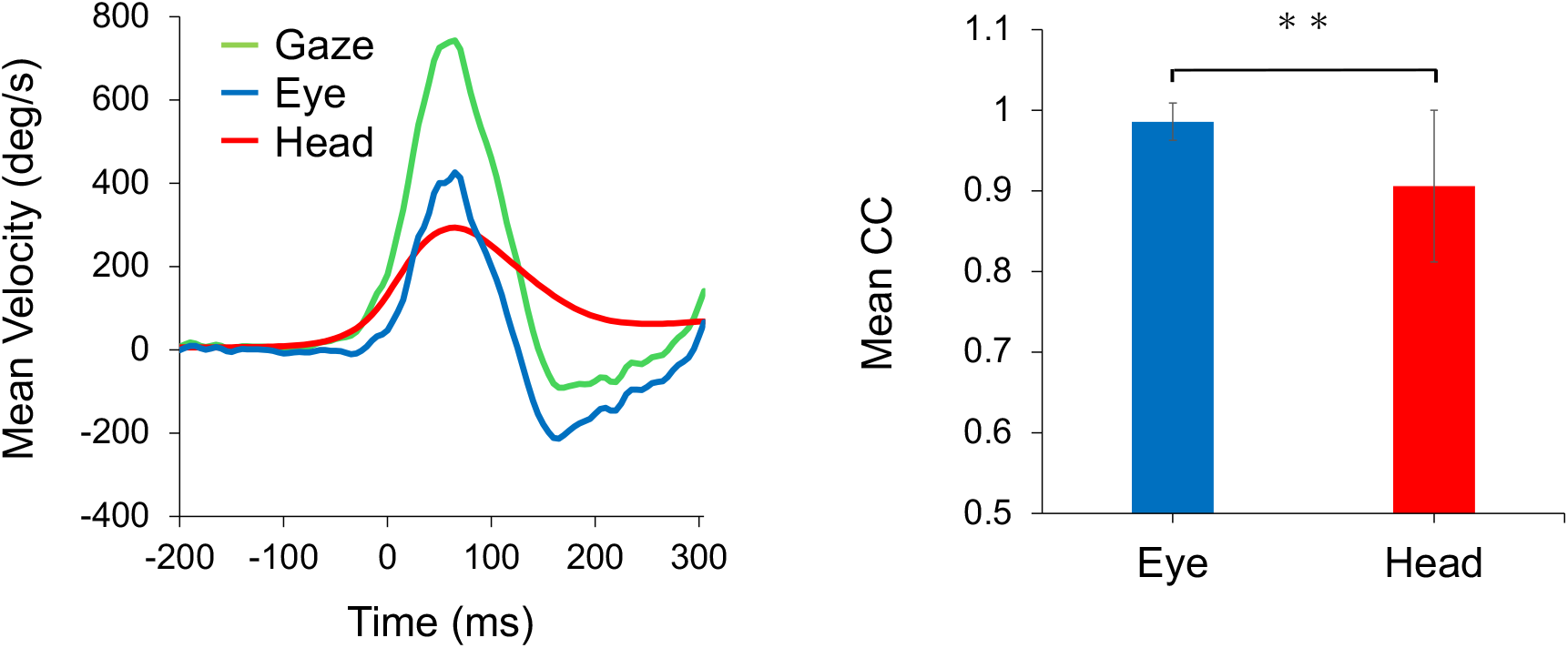
Representative mean angular-velocity waveforms. (A) Representative mean velocity traces for gaze (green), eyes (blue), and head (red). (B) Comparison of maximum cross-correlation (CC) for Gaze–Eye and Gaze–Head (**p < .001).

### 3.2. Relationships between eye and head velocities

Figure 5A shows the mean velocities of the eyes and head for each participant. Pearson’s correlation analysis revealed a negative correlation between the two variables (r = −.60, p = .005). As shown in Figure 5B, a positive correlation was observed for the velocity-control index—defined as mean velocity normalized by amplitude—for the eyes and head (r = .90, p < .001). These findings suggest that although individuals may differ in whether they emphasize the eyes or the head as the primary component of their gaze-shift strategy, the robustness of individual velocity-control competence is preserved across both strategies, as illustrated in Figure 5

**Figure 5.**
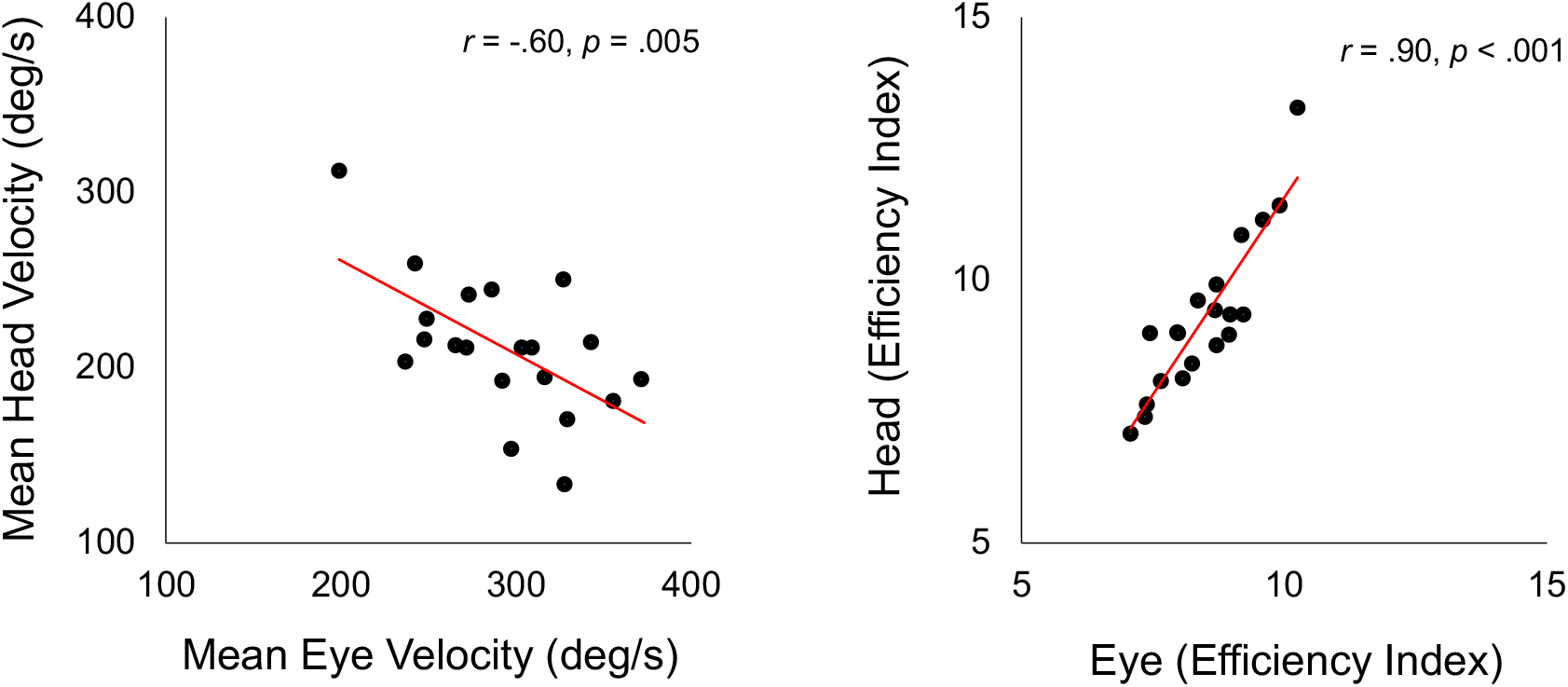
Correlation between eye and head velocities and control profiles. (A) Scatterplot of mean velocity (eye vs. head) across participants. (*p = .005). (B) Scatterplot of velocity-control profiles (mean velocity/amplitude) showing a positive correlation (**p < .001).

### 3.3. Eyes provide movement stability

Figure 6A shows the coefficients of variation (CV) of peak velocity for the eyes and head across participants. Paired t-tests revealed that the CV of peak velocity was significantly lower for the eyes than for the head (t(19) = 3.51, p = .002, d = 0.79). Figure 6B displays the CV of amplitude for the eyes and head. Paired t-tests further showed that the CV of amplitude was also significantly lower for the eyes (t(19) = 10.46, p < .001, d = 2.34). In contrast, no significant difference was found for the CV of mean velocity (t(19) = 0.76, p = .457, d = 0.17). These results indicate that the eyes exhibited a more stable movement profile compared with the head, as illustrated in Figure 6

**Figure 6.**
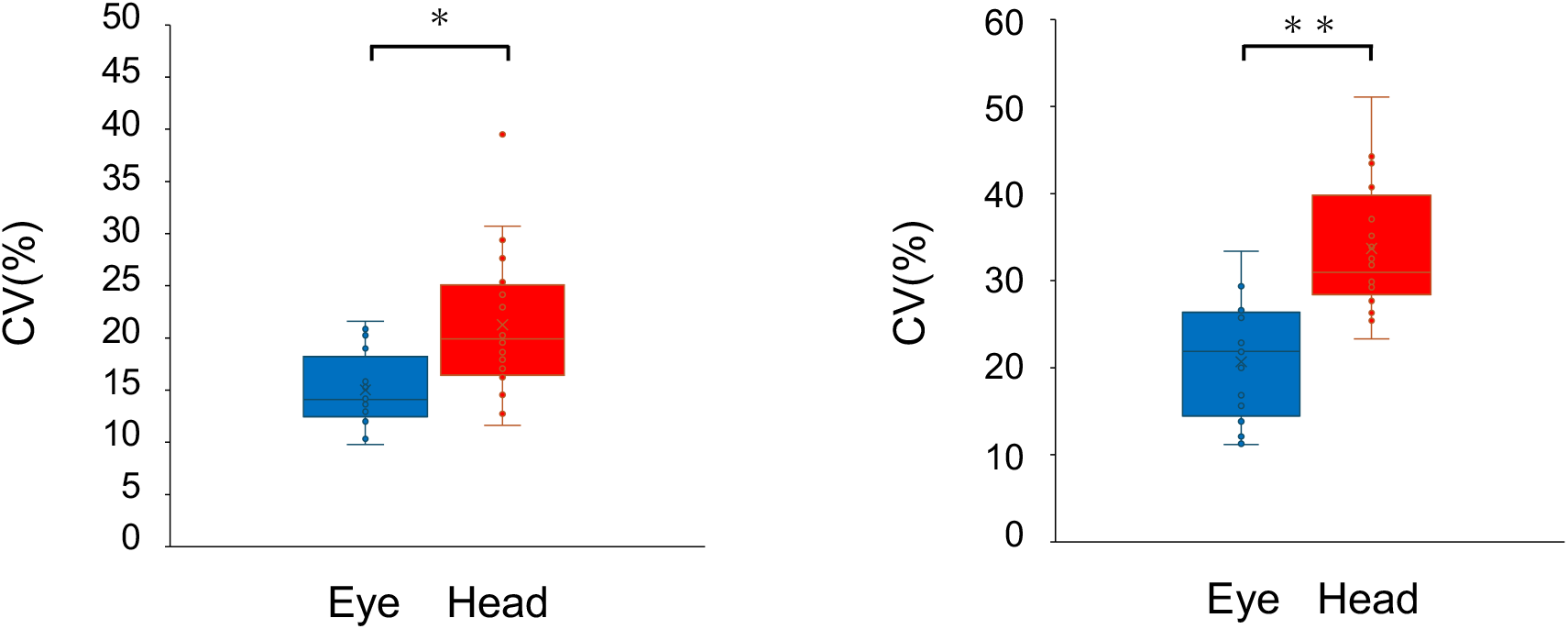
Coefficients of variation (CV) for eyes and head. (A) Peak-velocity CV lower for the eye (*p = .002). (B) Amplitude CV lower for the eye (**p < .001).

### 3.4. The head adapts to context

Figure 7 illustrates comparisons of peak velocity under two speed conditions (800 ms vs. 1600 ms) for the eye (A) and head (B) velocities. Paired t-tests revealed no significant difference between speed conditions for peak eye velocity (t(19) = 0.13, p = .898, d = 0.03). In contrast, Wilcoxon signed-rank tests showed that peak head velocity was significantly higher in the 800 ms condition (Z= 3.66, p < .001, r = .82). These findings indicate that while the eyes maintained consistent dynamics across speed conditions, the head adapted its movements to accommodate contextual changes. In other words, the head contributed more prominently during rapid gaze shifts under shorter time constraints, as illustrated in Figure 7

**Figure 7.**
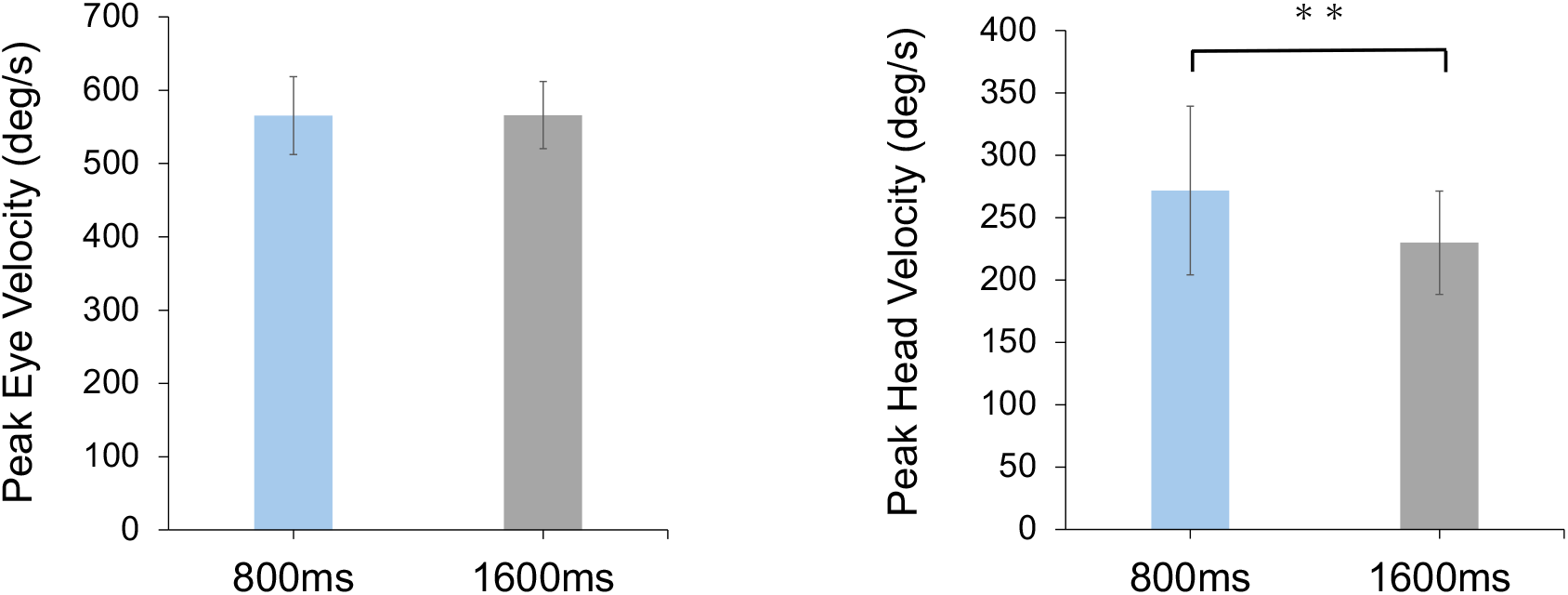
Peak velocity by time condition (800 ms vs. 1600 ms). (A) Peak eye velocity showed no significant difference between conditions. (B) Peak head velocity was significantly higher under time pressure (800 ms) compared with 1600 ms (**p < .001).

## 4 Discussion

### 4.1. Comparison between eye and head movements during gaze shifts

This study demonstrated that gaze shifts are predominantly led by the eyes rather than the head. Our results revealed that peak eye velocity and amplitude were significantly greater than those of the head, and the cross-correlation coefficient between gaze and eye was significantly higher than that between gaze and head. These results suggest that gaze shifts are more strongly driven by the eyes than by the head. Previous studies on scanning have primarily focused on the relationship between head movements and performance (Jordet et al., 2020; Berg et al., 2025). However, the contribution of the eye movement has not been sufficiently examined. The novelty of this study lies in the simultaneous measurement of head and eye movements, providing a quantitative characterization of the coordination structure of gaze shifts. Gaze is formed as the composite eye and head velocities, and cross-correlation serves as an index of how closely their temporal fluctuation patterns align. In this study, the eyes exhibited significantly higher peak velocity than the head, accompanied by sharper and faster fluctuations. Consequently, the rising slope and peak shape of the gaze waveform were more strongly influenced by the eye velocity, which likely explains why the gaze–eye cross-correlation coefficient was higher than that of gaze–head. In other words, the eyes represent a primary factor shaping the instantaneous form of gaze, supporting the interpretation that gaze shifts are strongly led by the eye velocity. By clarifying the leading role of the eye movement in scanning, the findings of this study provide an opportunity to reconsider practical scanning instruction, which has traditionally emphasized head turns. Future work may benefit from comprehensive measurement of both eyes and head, which could lead to more accurate assessments of situational decision-making ability.

### 4.2. Relationships between eye and head velocities

This study showed that participants differed in whether they generated velocity primarily through eye or head movements during gaze shifts. However, for both strategies, the velocity-control profile of the eye and head velocity (mean velocity relative to amplitude) was maintained in accordance with individual characteristics. Pearson’s correlation analyses revealed a negative correlation between the mean eye and head velocities, whereas the velocity-control profile showed a positive correlation. This coexistence of negative and positive correlations is consistent with previous research. Earlier studies have shown that as gaze amplitude increases, the contribution of the head motion also increases (Freedman et al., 2000), and that this contribution varies depending on an individual’s gaze-shift strategy (Fuller, 1992). The divergence observed in this study, where the contrast between participants with a high contribution of head movement to eye movements and those with the opposite pattern, aligns with these reports. Moreover, it is known that the superior colliculus (SC) generates common gaze-shift commands to both the eyes and head (Freedman & Sparks, 1997), suggesting that participants selected strategies suited to their motor characteristics under this shared command. As a result, the negative correlation between mean eye and head velocities, while the positive correlation in velocity-control profiles, can be interpreted as reflecting the expression of individual motor capacities. A previous study has reported substantial individual differences in the number of scans performed during actual matches, with disparities exceeding 100 instances (Aksum et al., 2021). The strategic diversity of gaze shifts identified in this study is consistent with such on-field individual differences and indicates the existence of distinct scanning styles among players.

Therefore, the findings of this study may directly inform the design of more individualized training programs. For example, head-dominant players may benefit from tasks that require rapid decision-making with stabilized eye control, whereas eye-dominant players may benefit from broader exploratory tasks that exploit the head’s range of motion, enabling training tailored to individual characteristics.

### 4.3. Eyes confer stability

This study demonstrated that the eye movements play a stabilizing role in gaze shifts. Our results revealed that the coefficients of variation (CV) for peak velocity and amplitude were significantly lower for the eye than for the head. The peak eye velocity of (565 ± 49 deg/s) was close to the saturation range of saccadic velocity (600–700 deg/s; Bahill et al., 1975), and the mean amplitude (34.8 ± 6.6°) was also near the upper limit of ocular range of motion (30–40°; Freedman, 2008). Thus, the eye movements operated near their physiological limits, leading to suppressed variability in velocity and amplitude and consequently to lower CV values. Moreover, because the eye movements are less affected by inertia and antagonist muscle activity (Robinson, 1981) and function as effectors that reliably execute superior colliculus (SC) commands, they exhibited more consistent movements compared with the head. The possibility that stability of gaze control contributes to the accuracy of decision-making has been noted in reviews of sports science (Hüttermann et al., 2018), and the present findings provide experimental support for this view. Ocular stability may underpin accurate decision-making, as evidenced by reports that elite soccer players’ superior visual function strongly correlates with executive functions related to decision-making, such as selective attention and cognitive flexibility (Knöllner et al., 2022). Future research should directly examine the relationship between ocular stability and actual decision-making accuracy, which may enable the application of these insights to training practice.

### 4.4. The head adapts to environmental demands

Our findings indicated that the head plays a role in flexibly adapting to environmental conditions. The results showed that peak head velocity was significantly higher in the 800 ms condition than in the 1600 ms condition, whereas peak eye velocity did not differ between conditions. This result reflects the fact that the eye movements, operating near their physiological limits, show limited variability in velocity and amplitude, whereas the head possesses a greater capacity for adjustment due to the cervical range of motion and muscular flexibility. Indeed, a previous study has shown that peak head velocity continues to increase with amplitude without saturation (Zangemeister et al., 1981), and the present findings are consistent with this. Thus, the findings suggest that under temporal constraints, the contribution of the head increases. Furthermore, it has been reported that temporal pressure alters exploratory behavior in sports contexts (Vater et al., 2020), which aligns with the present results. From a practical perspective, manipulating time constraints may emphasize the head’s contribution and facilitate training designs that promote broader exploration. For example, time-limited passing tasks based on the present experimental setup have been shown to elicit head-dominant exploration, suggesting potential applications for training.

### Limitations

This study has several limitations. First, although the task employed in this study simulated a practical soccer scenario, it did not fully replicate the complexity and dynamic interactions of actual environments. Therefore, direct conclusions regarding gaze behavior and decision-making in real game situations cannot be drawn. Second, the neural basis of eye–head coordination was addressed only through indirect inference; thus, future studies integrating neurophysiological measurements and computational modeling are needed for a more comprehensive understanding. Finally, because this study employed a cross-sectional design, it remains unclear to what extent the observed strategic characteristics can be modified through training. Future longitudinal and intervention studies are required to examine how training programs that emphasize the roles of eye and head movements influence decision-making ability and competitive performance.

## 5 Conclusion

This study is to empirically elucidate the structure of eye–head coordination during gaze shifts in soccer scanning. Our results showed that gaze shifts are driven primarily by eye movements rather than head movements. Furthermore, eye movements ensure gaze stability while head movements contribute to environmental adaptation. Furthermore, the identification of trade-offs in eye–head velocity relationships and the quantification of individual-specific velocity-control capacities represent a novel contribution that applies fundamental research insights to the sporting context. These advances enable a more comprehensive assessment of scanning beyond traditional head-turn indicators and provide insights directly relevant to evaluating situational decision-making ability and designing individualized training programs. Future research should compare across levels of expertise and contextual conditions, and further validate findings in practical situations, to comprehensively clarify the impact of eye–head coordination on decision-making and performance.

## 6 Conflict of Interest

The authors declare that the research was conducted in the absence of any commercial or financial relationships that could be construed as a potential conflict of interest.

## 7 Funding

This research was supported by JSPS KAKENHI Grant Number 23K24749

## 9. Data Availability Statement

The datasets generated for this study are available from the corresponding author on reasonable request.

## Notes

### Competing Interest Statement

The authors have declared no competing interest.

## Reference

Aksum, K. M., Brotangen, L., Bjørndal, C. T., Magnaguagno, L., and Jordet, G. (2021a). Scanning activity of elite football players in 11 vs. 11 match play: An eye-tracking analysis on the duration and visual information of scanning. PLOS ONE 16, e0244118. doi: 10.1371/journal.pone.0244118

Aksum, K. M., Pokolm, M., Bjørndal, C. T., Rein, R., Memmert, D., and Jordet, G. (2021b). Scanning activity in elite youth football players. Journal of Sports Sciences 39, 2401–2410. doi: 10.1080/02640414.2021.1935115

Bahill, A. T., Clark, M. R., and Stark, L. (1975). The main sequence, a tool for studying human eye movements. Mathematical Biosciences 24, 191–204. doi: 10.1016/0025-5564(75)90075-9

Berg, A., Malmsten, J., Lind, J., Mannix, H., Sjösten, L., Josefsson, A., et al. (2025). Scanning is associated with better performance in professional ice hockey. Journal of Sports Sciences 43, 145– 150. doi: 10.1080/02640414.2024.2433899

Chalkley, D., Shepherd, J. B., McGuckian, T. B., and Pepping, G.-J. (2018). Development and validation of a sensor-based algorithm for detecting the visual exploratory actions. IEEE Sensors Letters 2, 1–4. doi: 10.1109/LSENS.2018.2839703

Davids, K., Williams, A. M., and Williams, J. G. (eds.) (2005). Visual perception and action in sport. 1st Edn. London: Routledge doi: 10.4324/9780203979952

Faul, F., Erdfelder, E., Lang, A.-G., and Buchner, A. (2007). G*Power 3: A flexible statistical power analysis program for the social, behavioral, and biomedical sciences. Behavior Research Methods 39, 175–191. doi: 10.3758/BF03193146

Freedman, E. G. (2008). Coordination of the eyes and head during visual orienting. Experimental Brain Research 190, 369–387. doi: 10.1007/s00221-008-1504-8

Freedman, E. G., and Sparks, D. L. (1997). Activity of cells in the deeper layers of the superior colliculus of the rhesus monkey: evidence for a gaze displacement command. Journal of Neurophysiology 78, 1669–1690. doi: 10.1152/jn.1997.78.3.1669

Freedman, E. G., and Sparks, D. L. (2000). Coordination of the eyes and head: movement kinematics. Experimental Brain Research 131, 22–32. doi: 10.1007/s002219900296

Fuller, J.H. (1992). Head movement propensity. Experimental Brain Research 92, 152–164. doi: 10.1007/BF00230391

Gregori Grgič, R., Crespi, S. A., and de’Sperati, C. (2016). Assessing Self-Awareness through Gaze Agency. PLOS ONE 11, e0164682. doi: 10.1371/journal.pone.0164682

Hüttermann, S., Noël, B., and Memmert, D. (2018). Eye tracking in high-performance sports: Evaluation of its application in expert athletes. International Journal of Computer Science in Sport 17, 182–203. doi: 10.2478/ijcss-2018-0011

Jordet, G. (2005). Perceptual Training in Soccer: An Imagery Intervention Study with Elite Players. Journal of Applied Sport Psychology 17, 140–156. doi: 10.1080/10413200590932452

Jordet, G., Aksum, K. M., Pedersen, D. N., Walvekar, A., Trivedi, A., McCall, A., et al. (2020). Scanning, Contextual Factors, and Association With Performance in English Premier League Footballers: An Investigation Across a Season. Frontiers in Psychology 11, 553813. doi: 10.3389/fpsyg.2020.553813

Kishita, Y., Ueda, H., and Kashino, M. (2020). Eye and Head Movements of Elite Baseball Players in Real Batting. Frontiers in Sports and Active Living 2, 3. doi: 10.3389/fspor.2020.00003

Klatt, S., and Smeeton, N. J. (2022). Processing visual information in elite junior soccer players: Effects of chronological age and training experience on visual perception, attention, and decision making. European Journal of Sport Science 22, 600–609. doi: 10.1080/17461391.2021.1887366

Knöllner, A., Memmert, D., von Lehe, M., Jungilligens, J., and Scharfen, H.-E. (2022). Specific relations of visual skills and executive functions in elite soccer players. Frontiers in Psychology 13, 960092. doi: 10.3389/fpsyg.2022.960092

Kredel, R., Hernandez, J., Hossner, E.-J., and Zahno, S. (2023). Eye-tracking technology and the dynamics of natural gaze behavior in sports: an update 2016–2022. Frontiers in Psychology 14, 1130051. doi: 10.3389/fpsyg.2023.1130051

Manakhov, P., Sidenmark, L., Pfeuffer, K., and Gellersen, H. (2024). Filtering on the Go: Effect of Filters on Gaze Pointing Accuracy During Physical Locomotion in Extended Reality. IEEE Transactions on Visualization and Computer Graphics 30, 7234–7244. doi: 10.1109/TVCG.2024.3456153

Mann, D. T. Y., Williams, A. M., Ward, P., and Janelle, C. M. (2007). Perceptual-Cognitive Expertise in Sport: A Meta-Analysis. Journal of Applied Sport Psychology 29, 457–478. doi: 10.1123/jsep.29.4.457

McGuckian, T. B., Cole, M. H., Chalkley, D., Jordet, G., and Pepping, G.-J. (2019). Visual Exploration When Surrounded by Affordances: Frequency of Head Movements Is Predictive of Response Speed. Ecological Psychology 31, 30–48. doi: 10.1080/10407413.2018.1495548

McGuckian, T. B., Cole, M. H., Chalkley, D., Jordet, G., and Pepping, G.-J. (2020). Constraints on visual exploration of youth football players during 11v11 match-play: The influence of playing role, pitch position and phase of play. Journal of Sports Sciences 38, 658–668. doi: 10.1080/02640414.2020.1723375

Parker, P. R. L., Martins, D. M., Leonard, E. S. P., Casey, N. M., Sharp, S. L., Abe, E. T. T., et al. (2023). A dynamic sequence of visual processing initiated by gaze shifts. Nature Neuroscience 26, 2192–2202. doi: 10.1038/s41593-023-01481-7

Pokolm, M., Kirchhain, M., Müller, D., Jordet, G., and Memmert, D. (2023). Head movement direction in football - a field study on visual scanning activity during the UEFA-U17 and -U21 European Championship 2019. Journal of Sports Sciences 41, 695–705. doi: 10.1080/02640414.2023.2235160

Pokolm, M., Rein, R., Müller, D., Nopp, S., Kirchhain, M., Aksum, K. M., et al. (2022). Modeling Players’ Scanning Activity in Football. Journal of Sport and Exercise Psychology 44, 263–271. doi: 10.1123/jsep.2020-0299

Robinson, D. A. (1981). The use of control systems analysis in the neurophysiology of eye movements. Annual Review of Neuroscience 4, 463–503. doi: 10.1146/annurev.ne.04.030181.002335

Roca, A., Ford, P. R., McRobert, A. P., and Mark Williams, A. (2011). Identifying the processes underpinning anticipation and decision-making in a dynamic time-constrained task. Cognitive Processing 12, 301–310. doi: 10.1007/s10339-011-0392-1

Roca, A., Williams, A. M., and Ford, P. R. (2014). Capturing and testing perceptual-cognitive expertise: A comparison of stationary and movement response methods. Behavior Research Methods 46, 173–177. doi: 10.3758/s13428-013-0359-5

Silva, A. F., Conte, D., and Clemente, F. M. (2020). Decision-Making in Youth Team-Sports Players: A Systematic Review. International Journal of Environmental Research and Public Health 17, 3803. doi: 10.3390/ijerph17113803

Van Biemen, T., and Mann, D. L. (2024). How do referees visually explore? An in-situ examination of the referential head and eye movements of football referees. Journal of Sports Sciences 42, 1243– 1258. doi: 10.1080/02640414.2024.2387972

Van Biemen, T., Oudejans, R. R. D., Savelsbergh, G. J. P., Zwenk, F., and Mann, D. L. (2023). Into the eyes of the referee: a comparison of elite and sub-elite football referees’ on-field visual search behaviour when making foul judgements. International Journal of Sports Science and Coaching 18, 78–90. doi: 10.1177/17479541211069469

Vater, C., Williams, A. M., and Hossner, E.-J. (2020). What do we see out of the corner of our eye? The role of visual pivots and gaze anchors in sport. International Review of Sport and Exercise Psychology 13, 81–103. doi: 10.1080/1750984X.2019.1582082

Vítor De Assis, J., Costa, V., Casanova, F., Cardoso, F., and Teoldo, I. (2021). Visual search strategy and anticipation in tactical behavior of young soccer players. Science and Medicine in Football 5, 158–164. doi: 10.1080/24733938.2020.1823462

Zangemeister, W. H., Jones, A., and Stark, L. (1981). Dynamics of head movement trajectories: main sequence relationship. Experimental Neurology 71, 76–91. doi: 10.1016/0014-4886(81)90072-8

